## Supplementary figures and images for "High School Science Fair: What Students Say -- Mastery, Performance, and Self-Determination Theory"

### S1 Figure

S1 Figure. Distribution of students' comments in answer to the *Reason Why?* question year by year.

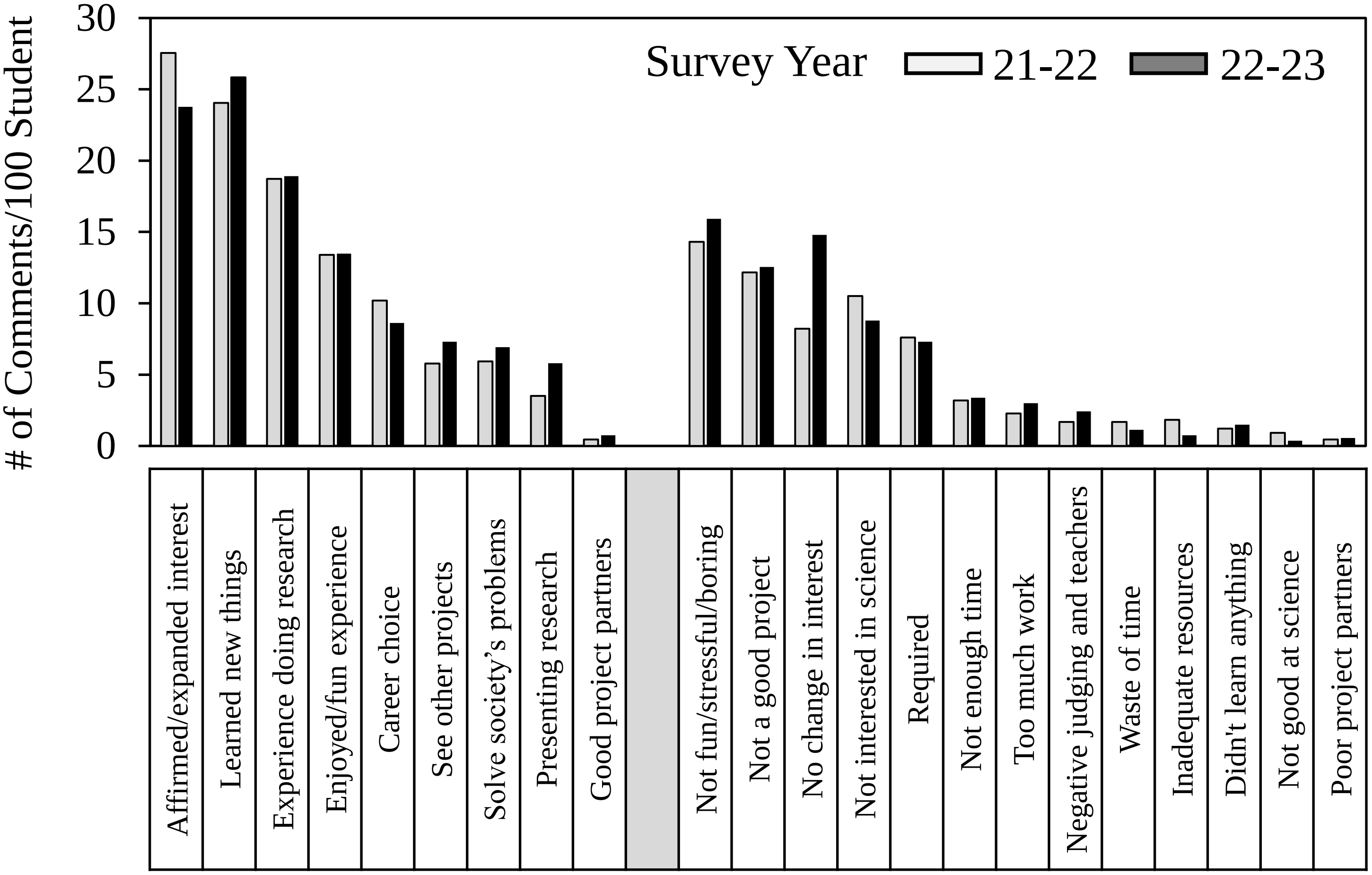
