## Supplementary material for "High School Science Fair: What Students Say -- Mastery, Performance, and Self-Determination Theory": S1 Survey

### High School Science Fair Survey

Scientists at UT Southwestern Medical Center and Southern Methodist University are interested in learning about student experiences with high school science fair, and we ask for your assistance. This survey, which consists of 20 questions, should only require about 10-15 minutes to complete. You may leave blank any question you prefer not to answer. The survey is anonymous; we only are interested in overall trends. It is important that you give honest replies to those questions that you do answer since the results may be used to influence science fair practices in the future. Your participation in the survey is voluntary; however, the more students who participate in the survey, the more statistically persuasive will be any conclusions. Thank you for your help.

---

Enter your survey access number... ______________________________

---

1. What grade are you in?
   - ◯ 9th
   - ◯ 10th
   - ◯ 11th
   - ◯ 12th

---

2. Location of high school?
   - ◯ Urban
   - ◯ Suburban
   - ◯ Rural

---

3. Gender?
   - ◯ Female
   - ◯ Male

---

4. Ethnicity most identified with?
   - ◯ Asian
   - ◯ Black or African American
   - ◯ Hispanic or Latino
   - ◯ Native Hawaiian or Other Pacific Islander
   - ◯ White
   - ◯ American Indian or Alaska Native
   - ◯ Other

---

Specify Other: ______________________________

---

5. During high school have you carried out science fair more than once?
   - ◯ Yes
   - ◯ No

---

If you carried out science fair more than once, then on subsequent questions use your most recent experience to answer.

---

6. In which science fair competitions did you compete this year?
   Check all that apply
   - ☐ School
   - ☐ District
   - ☐ Region
   - ☐ State

---

7. Was your science fair project Team or Individual?
   - ◯ Team
   - ◯ Individual

---

8. Was the science fair project required by your school?
   - ◯ Yes
   - ◯ No
   - ◯ No, but I did a science fair project to satisfy a school project requirement.

---

9. Do you think science fair projects should be optional or required? (This need not be for competition.)
   - ◯ Optional
   - ◯ Required

| | |
|---|---|
| 10. Do you think science fair projects for competition should be optional or required? | ◯ Optional<br>◯ Required |
| 11. Who helped you with your science fair project?<br>Check all that apply | ☐ 1. Parents<br>☐ 2. Siblings<br>☐ 3. Other family members (uncles, cousins, etc.)<br>☐ 4. Teachers<br>☐ 5. Other students<br>☐ 6. Scientists<br>☐ 7. A paid mentor<br>☐ 8. Articles on the Internet<br>☐ 9. Articles in books or magazines<br>☐ Other |
| Specify: | __________________________ |
| 12. What kind of help did you actually receive?<br>Check all that apply | ☐ 1. Being given the main idea<br>☐ 2. Development of the idea<br>☐ 3. Gathering background research information, or finding a research site or participants<br>☐ 4. Performing the experiments<br>☐ 5. Writing the report<br>☐ 6. Fine tuning the report after it is written<br>☐ 7. Designing the poster board and presentation<br>☐ 8. Producing charts or graphs<br>☐ 9. Coaching for the interview with judges<br>☐ 10. Copying the project from someone else<br>☐ Other |
| Specify: | __________________________ |
| 13. Did you get the kind of help you wanted from teachers? | ◯ Yes<br>◯ No |
| 14. Was there some kind of help that you would have liked but did not receive?<br>Specify: | __________________________ |
| 15. Did you get the amount of help you wanted from teachers? | ◯ Yes<br>◯ No |
| 16. What types of communications did you use in your science fair presentation?<br>Check all that apply. | ☐ 1. Written report<br>☐ 2. Literature review<br>☐ 3. Research notebook<br>☐ 4. Poster board preparation<br>☐ 5. PowerPoint presentation<br>☐ 6. Software to prepare tables, graphs, or images<br>☐ 7. Interview with the judges<br>☐ Other |
| Specify? | __________________________ |

| | |
|---|---|
| 17. What obstacles did you face?<br>Check all that apply | ☐ 1. Coming up with the main idea<br>☐ 2. Getting motivated to do the project<br>☐ 3. Becoming disappointed with the project<br>☐ 4. Limited resources<br>☐ 5. Limited knowledge<br>☐ 6. Limited skills<br>☐ 7. Limited cooperation<br>☐ 8. Getting organized<br>☐ 9. Time pressure<br>☐ 10. Not enough money<br>☐ 11. Results not as expected<br>☐ Other |
| Specify? | __________________________________ |
| 18. How did you overcome the obstacles you encountered?<br>Check all that apply | ☐ 1. Used someone else's main idea<br>☐ 2. Picked a familiar/interesting topic<br>☐ 3. Did more background research<br>☐ 4. Stopped working on the project for a while<br>☐ 5. Made a timeline to follow<br>☐ 6. Perseverance and self-discipline<br>☐ 7. Had someone else to keep me on track<br>☐ 8. Had someone else do the math<br>☐ 9. Changed the research plan<br>☐ 10. Collected more data<br>☐ 11. Had someone else collect the data<br>☐ 12. Used someone else's data<br>☐ 13. Made up the data<br>☐ 14. Changed the hypothesis to fit the data<br>☐ 15. Changed the data to fit the hypothesis<br>☐ Other |
| Specify? | __________________________________ |
| 19. Are you interested in a career in the sciences or engineering? | ◯ Yes<br>◯ No<br>◯ Not sure |
| 20. Did your science fair experience increase your interest in the sciences or engineering? | ◯ Yes<br>◯ No |
| 20A. Reason why? | __________________________________________ |
