## Supplementary material for "High School Science Fair: What Students Say -- Mastery, Performance, and Self-Determination Theory": S! Table

### S1 Table. Quantitative student responses to the SEF survey year by year. - Questions 1-9

| SEF Survey Questions and Answers | | % Students | |
|---|---|---|---|
| Questions | Answers | (21-22) 657* | (22-23) 534** |
| 1. What grade are you in? | 9 | 43.3 | 41.9 |
| | 10 | 30.2 | 32.0 |
| | 11 | 17.6 | 20.2 |
| | 12 | 8.7 | 5.8 |
| 2. Location of high school? | Urban | 20.8 | 27.0 |
| | Suburban | 72.8 | 66.9 |
| | Rural | 3.6 | 3.7 |
| 2A. Type of high school? (22-23 only) | Public | NA | 85.8 |
| | Private | NA | 7.1 |
| | Charter | NA | 6.2 |
| 3. Gender? | Female | 52.4 | 55.4 |
| | Male | 45.4 | 43.4 |
| 4. Ethnicity most identified with? | Asian | 30.4 | 29.0 |
| | Black | 7.9 | 9.2 |
| | Hispanic | 17.6 | 19.9 |
| | White | 38.9 | 36.5 |
| | Other* | 3.5 | 3.4 |
| | Specify | - | - |
| 5. During high school have you carried out science fair more than once? | Once | 67.3 | 67.4 |
| | > Once | 32.4 | 31.8 |
| 6. In which science fair competitions did you compete this year (could be more than one)? | School | 71.7 | 70.0 |
| | District | 32.4 | 27.0 |
| | Region | 24.5 | 30.3 |
| | State | 2.6 | 2.6 |
| 7. Was your science fair project team or individual? | Individual | 65.5 | 61.0 |
| | Team | 32.5 | 34.6 |
| 8. Was the science fair project required by your school? | Required | 64.9 | 59.2 |
| | Optional | 16.4 | 21.9 |
| | Project | 16.6 | 14.8 |
| 9. Do you think science fair projects should be optional or required? | Optional | 72.8 | 75.7 |
| | Required | 26.6 | 22.7 |
| 10. Do you think science fair projects for competition should be optional or required? | Optional | 87.4 | 85.4 |
| | Required | 12.2 | 12.9 |
| 11. Who helped you with your science fair project? | Parents | 51.5 | 53.0 |
| | Siblings | 14.0 | 11.0 |
| | Other family members | 5.6 | 6.6 |
| | Teachers | 54.9 | 54.9 |
| | Other students | 32.2 | 30.3 |
| | Scientists | 5.9 | 8.1 |
| | A paid mentor | 1.1 | 0.9 |
| | Articles on the Internet | 55.8 | 53.7 |
| | Articles in books or magazines | 18.5 | 18.0 |
| | Other | 3.0 | 6.6 |
| | Specify: | - | - |

S1 Table - Questions 10-13

| Questions | Answers | (21-22) 657* | (22-23) 534** |
|---|---|---|---|
| 10. Do you think science fair projects for competition should be optional or required? | Optional | 87.4 | 85.4 |
| | Required | 12.2 | 12.9 |
| 11. Who helped you with your science fair project? | Parents | 51.5 | 53.0 |
| | Siblings | 14.0 | 11.0 |
| | Other family members | 5.6 | 6.6 |
| | Teachers | 54.9 | 54.9 |
| | Other students | 32.2 | 30.3 |
| | Scientists | 5.9 | 8.1 |
| | A paid mentor | 1.1 | 0.9 |
| | Articles on the Internet | 55.8 | 53.7 |
| | Articles in books or magazines | 18.5 | 18.0 |
| | Other | 3.0 | 6.6 |
| | Specify: | - | - |
| 12. What kind of help did you actually receive? | Being given the main idea | 10.9 | 11.2 |
| | Development of the idea | 30.5 | 28.3 |
| | Gathering background research information, or finding a research site or participants | 46.8 | 41.0 |
| | Performing the experiments | 38.8 | 38.6 |
| | Writing the report | 13.8 | 11.0 |
| | Fine tuning the report after it is written | 28.7 | 29.8 |
| | Designing the poster board and presentation | 21.3 | 21.5 |
| | Producing charts or graphs | 17.5 | 16.3 |
| | Coaching for the interview with judges | 9.9 | 8.8 |
| | Copying the project from someone else | 0.8 | 0.2 |
| | Other | 5.9 | 9.2 |
| | Specify: | - | - |
| 13. Did you get the kind of help you wanted from teachers? | Yes | 84.3 | 80.1 |
| | No | 14.1 | 17.4 |

S1 Table - Questions 14-17

| Questions | Answers | (21-22) 657* | (22-23) 534** |
|---|---|---|---|
| 14. Was there some kind of help that you would have liked but did not receive? | Specify: | - | - |
| 15. Did you get the amount of help you wanted from teachers? | Yes | 81.5 | 76.8 |
| | No | 16.9 | 20.2 |
| 16. What types of communication and presentation skills did you use in your science fair project? | Written report | 51.5 | 57.9 |
| | Literature review | 15.3 | 19.5 |
| | Research notebook | 29.5 | 32.8 |
| | Poster board preparation | 62.2 | 78.5 |
| | Powerpoint presentation | 44.7 | 27.2 |
| | Software to prepare tables, graphs, or images | 46.0 | 52.1 |
| | Presentation to other students | 0.0 | 42.9 |
| | Interview with the judges | 43.0 | 50.4 |
| | Other | 2.4 | 0.9 |
| | Specify? | - | - |
| 17. What obstacles did you face? | Coming up with the main idea | 46.7 | 43.3 |
| | Getting motivated to do the project | 48.8 | 42.1 |
| | Becoming disappointed with the project | 24.0 | 26.8 |
| | Limited resources | 33.1 | 37.5 |
| | Limited knowledge | 26.0 | 28.5 |
| | Limited skills | 18.2 | 21.3 |
| | Limited cooperation | 12.8 | 11.0 |
| | Getting organized | 26.9 | 25.3 |
| | Time pressure | 60.5 | 59.2 |
| | Not enough money | 9.7 | 11.8 |
| | Results not as expected | 24.9 | 24.7 |
| | Other | 4.0 | 3.9 |
| | Specify? | - | - |

S1 Table - Questions 18-20A

| Questions | Answers | (21-22) 657* | (22-23) 534** |
|---|---|---|---|
| 18. How did you overcome the obstacles you encountered? | Used someone elses main idea | 1.5 | 1.7 |
| | Picked a familiar/interesting topic | 34.3 | 31.1 |
| | Did more background research | 48.8 | 50.2 |
| | Stopped working on the project for a while | 19.6 | 17.0 |
| | Made a timeline to follow | 27.1 | 26.6 |
| | Perseverance and self-discipline | 50.8 | 50.7 |
| | Had someone else to keep me on track | 15.0 | 16.7 |
| | Had someone else do the math | 0.3 | 0.2 |
| | Changed the research plan | 18.4 | 15.7 |
| | Collected more data | 28.0 | 23.6 |
| | Had someone else collect the data | 1.4 | 1.5 |
| | Used someone elses data | 0.6 | 0.2 |
| | Made up the data | 1.1 | 1.3 |
| | Changed the hypothesis to fit the data | 3.2 | 5.6 |
| | Changed the data to fit the hypothesis | 1.2 | 0.2 |
| | Other | 5.2 | 6.7 |
| | Specify? | - | - |
| 19. Are you interested in a career in the sciences or engineering? | Yes | 58.1 | 62.0 |
| | Not sure | 27.2 | 22.7 |
| | No | 14.7 | 14.8 |
| 20. Did your science fair experience increase your interest in the sciences or engineering? | Yes | 56.1 | 56.0 |
| | No | 43.8 | 44.0 |
| 20A. "Reason why? | - | - | - |
| *Students who answered 20A "Reason Why?" question (977 students total completed surveys) | | | |
| **Students who answered 20A "Reason Why?" question (813 students total completed surveys) | | | |
