## Supplementary material for "High School Science Fair: What Students Say -- Mastery, Performance, and Self-Determination Theory": S2 Table

### S2 Table. # comments/100 students in relationship to student demographics and SEF experiences

| Survey Question | | # of Comments/100 Student | | | | | | | | |
|---|---|---|---|---|---|---|---|---|---|---|
| SEF participation increased my interest in S&E | Yes | 42.4 | 30.1 | 21.9 | 13.8 | | 0.6 | 0.3 | 0.1 | 0.4 |
| | No | 2.5 | 4.4 | 2.7 | 4.0 | | 33.5 | 27.7 | 22.0 | 16.4 |
| Interest in a career in S&E | Yes | 30.2 | 23.3 | 13.6 | 12.2 | | 8.0 | 13.5 | 1.7 | 4.2 |
| | No | 7.4 | 6.8 | 8.0 | 3.4 | | 33.0 | 5.7 | 46.0 | 17.6 |
| | Unsure | 22.7 | 15.0 | 16.3 | 6.3 | | 21.0 | 13.7 | 7.3 | 9.3 |
| Science fair requirement | Optional | 33.0 | 27.7 | 17.4 | 11.2 | | 2.2 | 3.6 | 1.8 | 0.9 |
| | Required | 22.3 | 16.0 | 11.7 | 8.9 | | 18.4 | 15.1 | 12.4 | 10.0 |
| | Project | 25.0 | 21.3 | 18.1 | 10.6 | | 12.8 | 11.7 | 8.5 | 5.9 |
| Help from scientists | Yes | 28.0 | 32.9 | 15.9 | 19.5 | | 6.1 | 3.7 | 3.7 | 2.4 |
| | No | 24.6 | 17.8 | 13.3 | 8.7 | | 15.7 | 13.0 | 10.2 | 7.8 |
| Received coaching for interview | Yes | 33.9 | 30.4 | 16.1 | 16.1 | | 4.5 | 3.6 | 2.7 | 0.9 |
| | No | 23.9 | 17.6 | 13.2 | 8.8 | | 16.1 | 13.3 | 10.5 | 8.2 |
| Competition level | Beyond school | 33.3 | 24.1 | 16.7 | 13.5 | | 5.1 | 4.7 | 4.1 | 1.4 |
| | School only | 18.9 | 15.1 | 12.7 | 7.3 | | 21.4 | 18.1 | 13.2 | 12.0 |
| Grade in which students participated in SEFs | 9 | 24.2 | 17.5 | 15.1 | 9.8 | | 14.5 | 11.8 | 9.0 | 6.3 |
| | 10 | 22.0 | 16.5 | 10.3 | 8.1 | | 22.5 | 14.4 | 13.0 | 10.6 |
| | 11 | 28.6 | 23.7 | 12.1 | 9.8 | | 8.9 | 13.8 | 4.9 | 7.1 |
| | 12 | 30.7 | 23.9 | 20.5 | 12.5 | | 2.3 | 3.4 | 12.5 | 2.3 |
| Student ethnicity | Asian | 29.9 | 23.2 | 13.0 | 9.0 | | 9.6 | 9.0 | 5.9 | 5.6 |
| | Black | 24.8 | 15.8 | 11.9 | 3.0 | | 21.8 | 10.9 | 11.9 | 8.9 |
| | Hispanic | 30.2 | 13.5 | 14.9 | 13.1 | | 11.3 | 6.8 | 11.3 | 9.5 |
| | White | 19.5 | 18.4 | 12.9 | 10.0 | | 18.6 | 18.2 | 12.0 | 7.3 |
| Selected "Reason Why?" Categories | | Learned new things | Experience doing research | Enjoyed/fun experience | Career choice | | Not fun/ stressful/ boring | Not a good project | Not interested in science | Required |
